# Online Self-Report Data for Duchenne Muscular Dystrophy confirms natural history and can be used to assess for therapeutic benefits

**DOI:** 10.1101/012344

**Authors:** Richard T. Wang, Cheri Silverstein, J. Wes Ulm, Ivana Jankovic, Ascia Eskin, Ake Lu, Vanessa Rangel Miller, Rita M. Cantor, Ning Li, Robert Elashoff, Ann S. Martin, Holly Peay, Stanley F. Nelson

**Affiliations:** Department of Human Genetics, David Geffen School of Medicine, University of California, Los Angeles; Division of Cardiology, David Geffen School of Medicine, University of California, Los Angeles; PatientCrossroads, San Mateo, California; Department of Biomathematics, David Geffen School of Medicine, University of California, Los Angeles; Parent Project Muscular Dystrophy, Hackensack, NJ; Department of Pathology and Laboratory Medicine, David Geffen School of Medicine, University of California, Los Angeles

## Abstract

To assess the utility of online patient self-report outcomes in a rare disease, we attempted to observe the effects of corticosteroids in delaying age at fulltime wheelchair use in Duchenne muscular dystrophy (DMD) using data from 1,057 males from DuchenneConnect, an online registry. Data collected were compared to prior natural history data in regard to age at diagnosis, mutation spectrum, and age at loss of ambulation. Because registrants reported differences in steroid and other medication usage, as well as age and ambulation status, we could explore these data for correlations with age at loss of ambulation. Using multivariate analysis, current steroid usage was the most significant and largest independent predictor of improved wheelchair-free survival. Thus, these online self-report data were sufficient to retrospectively observe that current steroid use by patients with DMD is associated with a delay in loss of ambulation. Comparing commonly used steroid drugs, deflazacort prolonged ambulation longer than prednisone (median 14 years and 13 years, respectively). Further, use of Vitamin D and Coenzyme Q10, insurance status, and age at diagnosis after 4 years were also significant, but smaller, independent predictors of longer wheelchair-free survival. Nine other common supplements were also individually tested but had lower study power. This study demonstrates the utility of DuchenneConnect data to observe therapeutic differences, and highlights needs for improvement in quality and quantity of patient-report data, which may allow exploration of drug/therapeutic practice combinations impractical to study in clinical trial settings. Further, with the low barrier to participation, we anticipate substantial growth in the dataset in the coming years.

## INTRODUCTION

The study of rare diseases by traditional methods, such as clinical trials or natural history studies, is limited by the challenges and costs of recruiting an adequate sample. Self-report registry-based approaches have important potential advantages including the ability to cost-effectively grow the study sample, but at the risk of potential reductions in data quality [1]. One disease for which patient registries have the potential to impact our understanding of treatment is Duchenne muscular dystrophy (DMD; OMIM #310200). DMD is the most common form of progressive childhood-onset muscular dystrophy, affecting 1.3–1.8 per 10,000 males between the ages of 5–25 years in a recent US population survey [2]. Progressive muscle weakness leads to inability to ambulate in the second decade of life and ultimately death, usually from cardiorespiratory failure. DMD is caused by mutations in the X-linked *DMD* gene that typically result in the complete loss of expression of the dystrophin protein. Thus, the diagnosis of DMD is currently clearly and readily established in routine practice.

There are no curative therapies, but multiple independent clinical trials since 1974 [3] and meta-analyses firmly establish that chronic corticosteroids at a dose equivalent to 0.75/mg/kg/day of prednisone slow the progression of muscle weakness [4–10]. Recommendations from European Neuromuscular Centre (ENMC) and American Academy of Neurology (AAN) support the use of chronic steroids for DMD [11, 12]. However, the optimal dosing regimen and steroid type remain unclear [13], and physician/patient usage remains variable [14].

In addition to corticosteroids, there are a number of therapies and clinical trials in development based on the tremendous knowledge about DMD pathophysiology [15]. There are also numerous unproven alternative therapies variably used by patients/parents. Here, we examine outcome data derived from the DuchenneConnect Registry (www.duchenneconnect.org) to determine if they are characteristic of the overall DMD population and sufficient to observe the known therapeutic benefits of steroids. This provides support for exploring benefit/harm of other commonly used interventions. All inferences are based on delay age at loss of ambulation, a major milestone in the disease progression.

## METHODS

### Duchenne subject data source

We retrospectively analyzed and interpreted all data collected by DuchenneConnect between September 2007 and August 3^rd^, 2011 in order to assess the utility of known data. DuchenneConnect is an online registry that compiles patient-reported data on individuals with Duchenne or Becker muscular dystrophy [16]. Responses were collected via a web-based questionnaire, and additional data are obtained by staff review of genetic testing reports. The analysis presented here was derived from data entered at a single point in time for each subject: the date of last login. All data were censored at the age of each patient at which a given question was asked. Of the 2,285 DuchenneConnect registrants as of August 2011, we removed individuals without definitive DMD diagnosis, females or individuals who provided ambiguous sex information, and individuals who submitted data from low income countries as defined by the Organisation of Economic Co-operation and Development (OECD). This resulted in a final subset of 1,057 DMD individuals with outcome data (Fig. 1).

**Figure 1.**
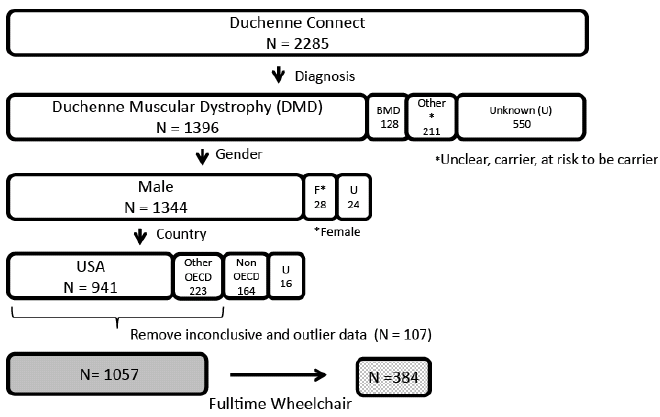
Study sample selection from Duchenne Connect Registry. Selection criteria included: an indicated diagnosis of DMD, male gender, and residence in one of the 34 Organization for Economic Co-operation and Development (OECD) countries. Of the 1,164 who met the criteria above, 107 were excluded due to concerns about data quality. 71 were excluded due to inconclusive data for the outcome variable: age at loss of ambulation. 29 subjects were excluded who first required a wheelchair more than 20 years ago, and 6 extreme outlier subjects were excluded, who reported ambulation at age 39 or above, as they are unlikely to have DMD. The final sample size was 1,057, of which 384 had reached the primary endpoint of loss of ambulation.

### Outcome variable

The outcome was age at fulltime wheelchair use (Age_WC_), a major functional milestone consistently reported by the majority of participants in the database and reliably recalled by participants. The registry includes several questions related to ambulation status. Subjects were considered wheelchair-dependent if they entered an age for the question “If a wheelchair is used all the time to move around, at what age did this become necessary?” Those without an entered age were considered ambulatory. If a participant entered a wheelchair age but also responded that he walked in response to the question “Are any devices used to help/assist with walking?” or did not select “Use a wheelchair fulltime” in response to “If strollers, wheelchairs, or scooters are used, then check all that apply below,” the patient was excluded. Similarly, participants were excluded if they left wheelchair age blank but selected “no, do not walk.”

### Treatment Variables

In the steroid-usage questionnaire, subjects could indicate whether they “never”, “previously”, or “currently” used steroids. Dosing frequency, but not exact dose, was solicited. For all other pharmacologic therapies, subjects could only select that the therapy was used. We defined a response as “use” and no response as “non-use,” an interpretation consistent with guidance provided by DuchenneConnect to registrants.

### Statistical Analyses

Statistical significance was accepted at p < 0.05. All analyses were performed in SAS, version 9.1 or 9.3 or R, version 2.10.1. No adjustments were made for multiple analysis of the primary outcome, as these data are considered exploratory.

#### Multivariate Modeling of Age_WC_

The primary analysis to assess the feasibility of using registry data was a comparison of Age_WC_ by corticosteroid use, controlled for other potentially confounding factors. Cox proportional hazards models were tested to identify the best predictive model for Age_WC_ and estimate hazard ratios for differing levels of steroid use (current, past, or never) [17]. Steroid use categories and covariates including (1) insurance status, (2) calcium use, (3) Vitamin D use, (4) the aggregate use of any of 9 other alternative therapies, (5) age of diagnosis, (6) United States residence, and (7) diagnosis confirmation by either genetic testing or biopsy, were entered into the initial model and backward stepwise selection with a threshold of p < 0.1 was employed to remove variables until the final model was identified. Log rank test sample sizes required for 80% power was computed at varying hazard ratios and treatment versus nontreatment proportions [18].

#### Comparison of Specific Steroid Regimens

To compare steroid regimens, Kaplan-Meier survival curves [19] were generated for the three reported categories of steroid use (current, past, or never) and the two categories of drug (deflazacort or prednisone) and compared using the log-rank test [20]. Those still ambulatory were censored at their age at the time of last login to DuchenneConnect.

#### Evaluation of Additional Therapeutic Agents on Age_WC_

The additional effects of the 11 most commonly-used supplemental therapies on Age_WC_ were explored in the subset of 633 individuals who reported current steroid use. We compared Kaplan-Meier curves with and without each agent using log-rank test. Cox proportional hazards regression with robust sandwich estimation and backward stepwise selection with a threshold of p < 0.1 were used to test the association between the 11 variables and Age_WC,_ including steroid type and dosing frequency.

### Standard protocol approvals, registrations, and patient consents

Participant consent for research was obtained at the time of DuchenneConnect registration. All data were downloaded from DuchenneConnect without personal identifiers. This study was deemed exempt from review by the University of California Los Angeles Institutional Review Board.

## RESULTS

### General characteristics of DuchenneConnect

The DuchenneConnect Registry sample analyzed here is comparable to other large natural history studies for DMD. Subjects in our sample reported years of birth between 1976 and 2010 with a median year of birth of 2000. Age of diagnosis is symmetrically distributed with a mean of 4.0 years (±2.3). Among the subset of 768 who indicated that they did not have a known family history of DMD, the mean age of diagnosis was 4.2 years, similar to published data [21]. The distribution of Age_WC_ is right-skewed with a median of 10 years (IQR 9–12) and a mean age of 10.5 years (±2.5) (Fig. 2) among the 384 subjects who already report fulltime wheelchair use. In our cohort, 17 of 1057 were reported to have died, and, of these individuals, the median age at death was 20 years (IQR 16–22). Genetic testing was performed in 94.7% of the overall sample. Genetic reports were provided for 688 subjects: 73% had an exon(s) deletion and another 3% had a smaller deletion. Approximately 10% had exon(s) duplication, 8% had a nonsense mutation, and about 3% had a splice-site mutation. Small duplications, insertions, combined frameshift insertion and deletion mutations, and either “other” or “no result provided” accounted for the remaining 3%. The distribution of mutations was similar among the 384 subjects who had already progressed to fulltime wheelchair use, and is consistent with other large registries of DMD mutations [22, 23].

**Figure 2.**
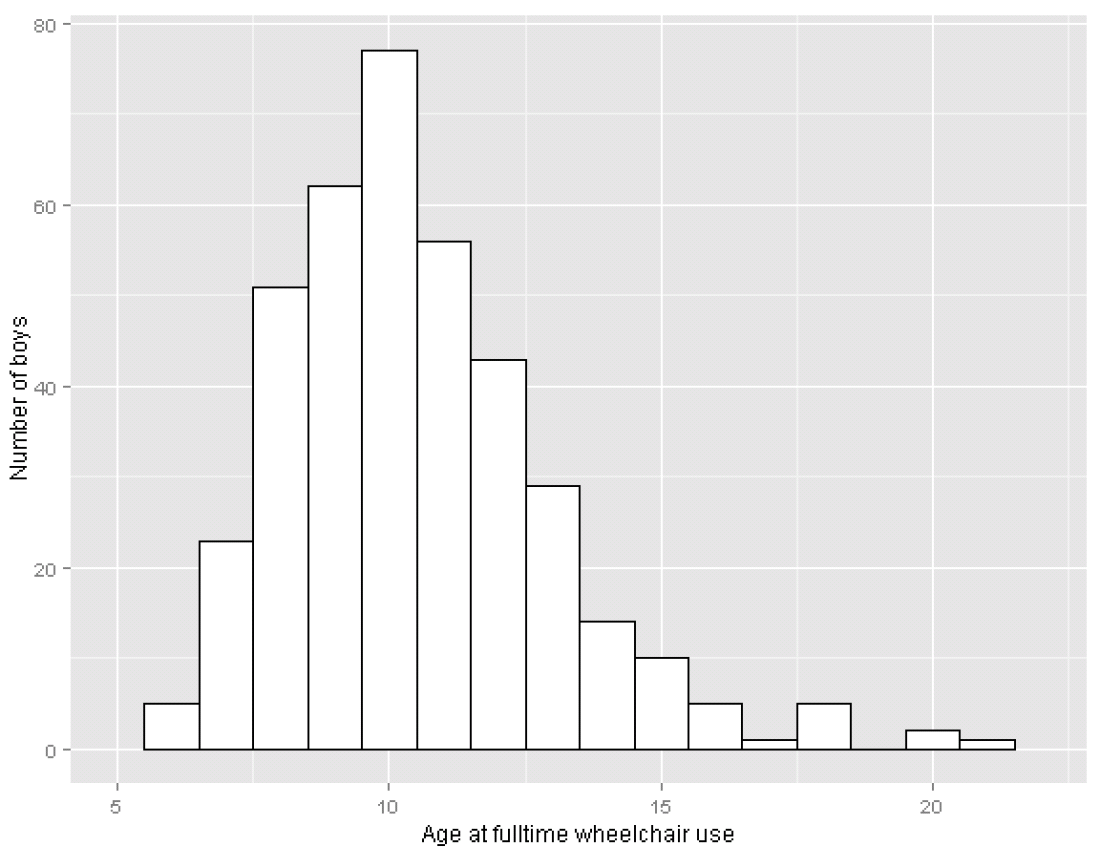
Age at loss of ambulation among those using a wheelchair fulltime. Histogram of age at loss of ambulation for the 384 males who reached that endpoint.

### Corticosteroid use and wheelchair-free survival in DuchenneConnect

Steroids were the most commonly used medication, consistent with their status as the only reliably identified medication to slow the progression of muscle weakness. We thus could compare those registrants currently using any dose of steroids (n = 633) with those individuals who report never having used steroids (n = 280) and previous users of steroids (n = 144). These data effectively allow us to determine if the online self report registry data can demonstrate the known therapeutic benefit of steroids, predicted here to be associated with prolonged ambulation. Consistent with this expectation, current steroid use was highly predictive of increased age at wheelchair use. Because we have many variables reported in DuchenneConnect, we can also explore the effect of other therapies reported. Multivariate analyses controlling for reported potential confounders found Vitamin D use, having insurance, and age at diagnosis greater than 4 years to be independent predictors of Age_WC_, along with current steroid use (Table 1). Steroid use was the most significant predictor (p < 0.0001).

**Table 1.** Cox proportional hazards model of time to fulltime wheelchair use by steroid use category (N = 928). Variables with p > 0.1 were eliminated from the model.

| Covariate | HR (95% CI) | P |
| --- | --- | --- |
| Current steroid use | 0.35 (0.28-0.43) | < 0.0001 |
| Has Insurance | 0.44 (0.23-0.84) | 0.01 |
| Vitamin D use | 0.66 (0.51-0.85) | 0.001 |
| Age at diagnosis > 4 | 0.78 (0.64-0.95) | 0.01 |

HR: Hazard Ratio; CI: Confidence Interval; P: P-value

### Comparison of Steroid Regimens in DuchenneConnect

Since different steroid types and regimens were reported, we could explore differences in relation to Age_WC_ prediction. To compare different steroid use categories, the Age_WC_ among subjects with outcome data available (the 633 subjects currently on steroids, 144 subjects previously on steroids, and 280 subjects who had never taken steroids) were analyzed using Kaplan-Meier analysis (Figure 3A). Current steroid use was significantly associated with longer wheelchair-free survival when compared with “past” and “never” steroid users (Fig. 3A; p < 0.0001). Among those who had progressed, median Age_WC_ was extended approximately 3 years, from less than 10 years for ‘never’ and ‘past’ to 13 years for the ‘current’ steroid use group. The effect of dose sizes could not be tested as that information is not currently captured. In DuchenneConnect, deflazacort was used by approximately 58% of those taking steroids, and, by Kaplan-Meier analysis, prolonged ambulation to a median of 14 years compared to prednisone which delayed wheelchair free survival to a median of 13 years (p = 0.0013, Fig. 3B). Consistent with clinic surveys showing that daily dosing is most common, approximately 85% of current steroid users in DuchenneConnect report taking steroids daily [14, 24]; however, less-than-daily dosing was more common among prednisone users (25% vs 9% for deflazacort, p < 0.0001). In both the prednisone and deflazacort subgroups, there was no significant difference in Age_WC_ between the daily and less-than-daily regimens (Fig. 3C and 3D).

**Figure 3.**
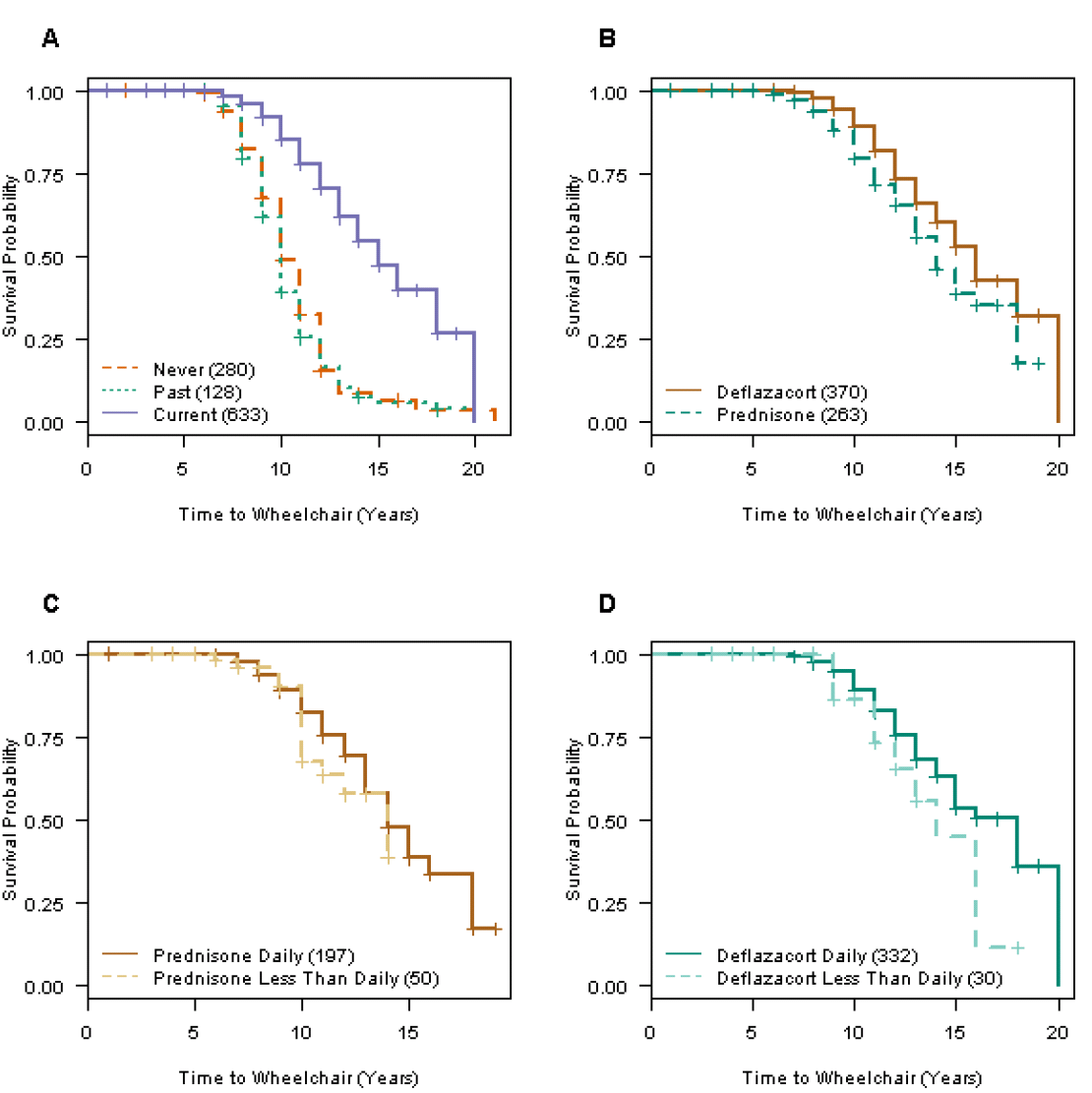
Effect of steroid treatments on wheelchair-free survival. Kaplan-Meier curves of wheelchair-free survival stratified by (A) steroid use category (p <0.0001, log rank test), (B) deflazacort versus prednisone (p=0.0013), (C) prednisone dosing frequency, and (D) deflazacort dosing frequency. All less-than-daily regimens are grouped together in C and D.

### Alternative supplements and wheelchair-free survival

Given that the documented effect of steroids in slowing disease progression could also be inferred from the available DuchenneConnect data, we sought to determine whether we could observe a potential additional effect from the use of any supplements reported. There were 11 different supplements used by at least 20 individuals (listed in Table 2), and for most, there were no available DMD studies. To test for a potential therapeutic benefit, for each supplement, we compared the subset of individuals taking that supplement who were also current steroid users to the 303 individuals who were current steroid users and did not report the use of any of the 11 supplements. Since loss of ambulation in this cohort occurred at a mean age of 12, we used Kaplan-Meier estimates to compare the probability of ambulating through the 12^th^ year of life for each combination (Table 2). In these analyses, only Vitamin D and Coenzyme Q10 have an additive effect and are significantly associated with longer ambulation. Power analysis at varying hazard ratios indicated that our sample sizes were generally too small to find any possible effect (Table 3).

**Table 2.** Kaplan-Meier estimates of time to loss of ambulation. P values represent the log-rank test of current steroid use plus each supplemental therapy compared to current steroids alone. Values for probability of walking through age 12 are derived from Kaplan-Meier lifetable probability estimates.

| Supplement | Probability of walking through age 12 | N | P value |
| --- | --- | --- | --- |
| Current steroids alone | 0.54 | 303 | NA |
| +Vitamin D | 0.72 | 246 | 0.004 |
| +Calcium | 0.68 | 207 | 0.07 |
| +Coenzyme Q10 | 0.74 | 155 | 0.007 |
| +Vitamin C | 0.57 | 41 | 0.43 |
| +Vitamin E | 0.73 | 28 | 0.06 |
| +Protandim | 0.92 | 44 | 0.22 |
| +Creatine Monohydrate | 0.75 | 33 | 0.10 |
| +Magnesium | 0.63 | 21 | 0.75 |
| +Melatonin | 0.58 | 21 | 0.39 |
| +L-Argenine | 0.94 | 21 | 0.22 |
| +Green Tea Extract | 0.75 | 17 | 0.5 |

**Table 3.** Sample sizes required for 80% power to detect effect. Each cell denotes the total sample size required. For example, if HR is 1.5 and 20% of individuals are treated, a study requires a total of 459 patients (367 untreated, 92 treated).

|  | **Proportion of individuals assigned to treatment** |  |  |  |  |
| --- | --- | --- | --- | --- | --- |
| **HR** | **0.1** | **0.2** | **0.3** | **0.4** | **0.5** |
| **1.1** | 13029 | 7296 | 5534 | 4821 | 4608 |
| **1.2** | 3701 | 2063 | 1558 | 1351 | 1286 |
| **1.3** | 1856 | 1030 | 774 | 668 | 633 |
| **1.4** | 1170 | 646 | 484 | 416 | 393 |
| **1.5** | 835 | 459 | 342 | 293 | 275 |
| **2.0** | 335 | 180 | 131 | 110 | 102 |
| **3.0** | 171 | 88 | 62 | 50 | 45 |

We performed multivariate Cox proportional hazard regression with backward stepwise selection with a threshold of p < 0.1 to determine whether the effects of Vitamin D and Coenzyme Q10 were supported and independent of other variables. In the sample of 633 subjects, use of deflazacort (HR = 0.68, 95% CI = 0.51–0.92), Vitamin D (HR = 0.75, 95% CI 0.55–1.03), and Coenzyme Q10 (HR = 0.68, 95% CI = 0.47–0.98) remained in the model as independent significant predictors. Thus, Vitamin D use and Coenzyme Q10 use appear to be independently and significantly associated with Age_WC_.

### Vitamin D and bone fracture risk

The observation of a beneficial effect of Vitamin D on loss of ambulation is provocative in DMD without a clear rationale from the available data, so we considered several potential explanations. One possibility is that Vitamin D usage lessened steroid-induced osteopenia, which is universally observed in DMD, and that this in turn lessened the risk of leg fracture and subsequent wheelchair use due to fracture. To perform an exploratory assessment of whether the association between Vitamin D usage and higher Age_WC_ was related to a difference in bone fractures, we compared the incidence of patient-reported leg and spinal fractures. Using a test of proportions, we found no difference in bone fracture rates in those reporting Vitamin D usage relative to those not reporting usage (p = 0.56) (data not shown).

## DISCUSSION

Online registries such as DuchenneConnect allow rapid and widespread accumulation and dissemination of data, inform researchers and practitioners about natural history, and facilitate recruitment into clinical trials [25]. In rare diseases, the low barrier to entry provides a major advantage allowing collection of data from a geographically broad area. Information can be gathered relatively inexpensively in electronic format, questions can be updated by scientists and clinicians, and new data can be generated to stimulate novel alternative means to solicit patient input. This set of tools opens the potential to more quickly explore off label treatment and patient-initiated treatment interactions that are harder to investigate in controlled clinical trials, and is thus a useful adjunct in understanding DMD and treatment response. Potential limitations are apparent, however. We had 2285 patients in the registry, but enforcement of quality control metrics (definitive DMD diagnosis, countries with high standards of advanced health care) censored over half the participants (Fig. 1). Because these data censoring operations selected individuals for whom data was entered consistently, the resulting final and interpretable data set likely enriches for families with better knowledge of advocacy guidelines, increased healthcare access, greater financial resources and internet savviness. In addition, geographic or national preferences for treatments of Duchenne muscular dystrophy may affect our interpretation of these results. Inherent biases in collection and analysis may affect the applicability of our findings in other populations in conjunction with more typical problems in online data collection such as inaccuracy of reporting and misclassification.

Self-report registry data are also limited due to the questions asked. Compared to randomized controlled trials, the major deficit of an online self-report registry is the possibility of unknown systematic bias among the respondents (e.g. family income allowing greater intensity of physical therapy, which may then be associated with steroid treatment or steroid type choice). Further, the prescription of steroid type and regimen may reflect physician bias in individual practice that is correlated with other unrecorded therapeutic procedures. As DuchenneConnect reaches a global audience, language barriers may also lead to some incorrect or misinterpreted responses, although we did not observe any strong differences between US and non-US portions of the sample set.

Only 13% of our sample had reported current or prior participation in any clinical research. Thus, this study substantially increased the percentage of individuals with DMD who have participated—notably with low participation burden. Based on the MD STARnet estimated population prevalence of DMD, there are about 6,000-8,000 DMD patients between the ages of 5–25 years in the US [21]. The 890 US patients in our analyzed sample therefore are a substantial fraction of the overall US DMD population, and compose a large single cohort of DMD with outcome measures. However, even with this substantial participation, there remains tremendous potential to grow this online sample.

Overall, the data from DuchenneConnect matched well with literature reports for mutation spectrum, age at diagnosis, and age at loss of ambulation [26, 27]. Early natural history studies, prior to the relatively common practice of chronic steroid usage, reported an average age at loss of ambulation of 9.5 years [28, 29]. Correspondingly, we found a median age at loss of ambulation of 9.9 years (IQR 8–11 ) in the 54 boys in our analysis who had never taken steroids or any supplements.

Steroids have been repeatedly documented to slow the skeletal muscle disease progression of DMD [3-10], but the use of steroids is not universal. The observed variation in clinical practice permitted us to assess the therapeutic benefit from the self-report data, as 40% of DMD patients do not report current treatment with steroids in this cohort. We note that the usage of steroids is actually higher than recent surveys. For instance, the MD STARnet study on the prevalence of corticosteroid use in patients with Duchenne and Becker muscular dystrophy found that 47.1% of 5- to 9-year-olds used corticosteroids in 2005, up from 35.3% in 1991 [30]. In our sample of 1057 DuchenneConnect Registry participants, 60% of the total sample reported current use with 70% use in those over 7 years of age, suggesting that there may be benefit from further increases in steroid usage in DMD. Although there is no clear current guidance on the use of corticosteroids after loss of ambulation, it is interesting to note that over 40% of patients who had progressed to using a wheelchair fulltime in our sample reported current steroid use.

From our analysis, DuchenneConnect data strongly support the therapeutic benefit of steroids to delay age at loss of ambulation, consistent with the prior clinical trial literature. An important caveat to our analysis is that specific dosing of steroids is not captured in DuchenneConnect, although clinical surveys indicate the most commonly prescribed doses are 0.75mg/kg/day of prednisone daily, or an equivalent daily dose of deflazacort, 0.9mg/kg/day [14, 24].. Future versions of the registry will solicit exact dose. Although not approved for use in DMD in the US, deflazacort is commonly used by the US participants. This usage is likely driven largely by reports of decreased side effects from deflazacort compared to prednisone, including reduced bone mineral loss, less weight gain, and good evidence of a therapeutic benefit [31, 32]. Interestingly, we found that those reporting usage of deflazacort instead of prednisone had a significant delay in age at loss of ambulation. Thus, these data support the continued use of deflazacort. However, the association of greater therapeutic benefit from deflazacort must be interpreted with some caution as no correction was made for multiple testing and no data were provided on the exact regimen, age at initiation, or reason for choosing one steroid type or for using a given supplement/vitamin.

Further, while we did not observe a significant difference between steroids dosed on a daily basis versus reduced dosing frequencies in the effect on age at loss of ambulation, the relatively small numbers of patients on less-than-daily regimens limited the power of our analyses. Nevertheless, these data raise the possibility that less-than-daily deflazacort may provide the optimal balance of muscle protection and reduced side effects. A recent study found that daily prednisolone significantly extended time to loss of ambulation and increased adverse side effects compared to less-than-daily prednisolone, but the effects of deflazacort were not studied [33]. Ongoing and future randomized trials that address optimal steroid regimen, such as the currently-recruiting FOR-DMD clinical trial [13], may help resolve these issues.

The confirmation of the therapeutic benefit of steroid usage demonstrates the potential utility of DuchenneConnect data for exploratory purposes of other potential therapeutic agents. We also identified Coenzyme Q10 and Vitamin D as therapeutic interventions that appear to benefit DMD patients, especially those on steroids. We observed an independent association of Vitamin D usage with higher Age_WC_, which appeared to be unrelated to bone fracture risk. Coenzyme Q10 is associated with prolonged ambulation in those DMD patients also taking steroids, which supports a recent smaller scale six month clinical trial of Coenzyme Q10 in DMD [34]. It is unclear if there is any relationship between the potential therapeutic benefit of Vitamin D and the reported observation of a relatively high frequency of Vitamin D deficiency in boys with DMD [35], but the data from DuchenneConnect suggest that broader usage of Vitamin D supplementation might be worth exploring as a therapeutic adjunct.

In our analysis of supplements other than Vitamin D and Coenzyme Q10, several of the supplements demonstrated longer wheelchair-free survival, however there were insufficient observations to achieve statistical significance because of the small number of subjects reporting usage of many of the supplements. With more registry participants, there may be an opportunity to confirm the Vitamin D and Coenzyme Q10 findings in a new set of patients, increase power to detect potential effects from other supplemental therapies, and investigate treatment interactions.

## CONCLUSION

The data presented here establish DuchenneConnect as a useful framework for further growth in online patient self-report data that is valuable for therapeutic interpretations regarding DMD, and support the more general use of online self-report data for therapeutic interpretation in rare disease. In five years, DuchenneConnect has accumulated the largest DMD cohort with age at full time wheelchair use and treatment information published to date, likely because of the low barrier to participation. Further, with many current registrants passing major milestones, like loss of ambulation, and the ongoing inclusion of new registrants, DuchenneConnect is poised to provide an ongoing and important longitudinal resource for observation of current DMD clinical care to support observations from clinical networks, clinical trial networks, or other natural history data. Thus, continued collection of data from web-registered participants promises to provide important natural history, clinical practice, and treatment response data for DMD. This study represents a successful, mutually beneficial collaboration between academic investigators and a patient self-report registry supported by an advocacy organization. Consistent with recent moves to embrace patient generated data, this study and the ongoing collaboration are informing the efficient use of patient-report data to describe patient outcomes and guide care provision.

## ACKNOWLEDGEMENTS

We thank the families who provided their data in DuchenneConnect. CAS was supported by T32-HL007895 and the UCLA STAR Program. IJ was supported by the UCLA Medical Scientist Training Program. Analytical work was supported by the Center for Duchenne Muscular Dystrophy at UCLA through funding by the National Institute of Arthritis, Musculoskeletal, and Skin Disorders (P30 AR057230) and the NIH/National Center for Advancing Translational Science (NCATS) UCLA CTSI UL1TR000124. The funders had no role in study design, data collection and analysis, decision to publish, or preparation of the manuscript.

